# A highly endemic area of *Echinococcus multilocularis* identified through a comparative re-assessment of prevalence in red fox (*Vulpes vulpes*), Alto Adige (Italy: 2019-2020)

**DOI:** 10.1101/2021.10.22.465400

**Authors:** Federica Obber, Roberto Celva, Graziana Da Rold, Karin Trevisiol, Silvia Ravagnan, Patrizia Danesi, Lucia Cenni, Chiara Rossi, Paola Bonato, Katia Capello, Heidi C. Hauffe, Alessandro Massolo, Rudi Cassini, Valentina Benvenuti, Andreas Agreiter, Davide Righetti, Marco Ianniello, Debora Dellamaria, Gioia Capelli, Carlo V. Citterio

## Abstract

**Background:** Surveillance of *E. multilocularis* at the edge of its range is hindered by fragmented distributional patterns and low prevalence and burden in definitive hosts. Thus, tests with adequate levels of sensitivity are especially important for discriminating between infected and non-infected areas.

**Aim:** We reassessed the prevalence of *E. multilocularis* at the southern border of its distribution in Alto Adige (Italy), to improve surveillance in wildlife and provide more accurate estimates of exposure risk.

**Methods:** We compared results from the diagnostic test currently implemented for surveillance (based on Coproscopy+Multiplex PCR - CMPCR), against a real-time quantitative PCR (qPCR) for 235 fox faeces collected in 2019-2020. The performances of the two tests were estimated using a scraping technique (SFCT) as the gold standard applied to the small intestines of a subsample (n=123) of the same hosts. True prevalence was calculated and sample size required by each faecal test for the detection of the parasite was then estimated.

**Results:** True prevalence of *E. multilocularis* in foxes (14.3%) was definitely higher than reported in the last decade (never >5% from 2012 to 2018). The qPCR also had a higher sensitivity (83%) compared to CMPCR (21%). Agreement with the gold standard was far higher for qPCR (0.816) than CMPCR (0.298) as well, determining a smaller sample size required to detect the disease.

**Conclusions:** Alto Adige should be considered a highly endemic area. Surveillance at the edges of *E. multilocularis* distribution should adopt qPCR diagnostics on definitive hosts on a small geographic scale.

**AUTHOR SUMMARY:** *Echinococcus multilocularis* is an intestinal flatworm, whose adult stage in Europe is harboured mainly by the red fox (*Vulpes vulpes*), which spreads parasite’s eggs by faeces. This parasite is the agent of a severe and potentially lethal zoonosis, the alveolar echinococcosis, affecting humans after accidental ingestion of parasite’s eggs. In the Italian Alpine area, which represents the southernmost border of *E. multilocularis* European range, surveillance is hindered by a fragmented distributional pattern, where presence in foxes has been consistently reported only in few isolated foci in Alto Adige (Bolzano province – Italy) of low prevalence. In order to improve the efficiency of monitoring efforts, we tested the performances of two diagnostic protocols on fox faeces (sedimentation, filtration, counting technique followed by standard PCR, and whole stool real-time PCR) against a benchmark technique on fox intestines (scraping, filtration, counting technique - considered as the gold standard). This allowed not only to determine qPCR as a far more sensitive and sample-efficient diagnostic tool for *E. multilocularis* detection in marginally affected areas, but also to re-assess its prevalence in Alto Adige, which should be considered a highly endemic area. Consequent actions in the field and modifications in the surveillance strategy should be therefore considered.

## INTRODUCTION

*Echinococcus* spp. (Cestoda, Cyclophyllidea, Taeniidae) are small intestinal tapeworms causing zoonoses of public health importance worldwide. In the European Union (EU), it is mandatory to report the detection of these pathogens to national authorities, and their surveillance, prevention and control are closely regulated for pets, livestock and wildlife.

Collection of relevant metadata is also highly recommended by both the Directive EU 2003/99/EC (monitoring of zoonoses and zoonotic agents) and the Regulation EU 2016/429 (‘Animal Health Law’). Among these, *Echinococcus multilocularis* has a complex life-history. The sylvatic cycle depends on a predator-prey system, where the adult (strobilar) stage of the parasite is carried by wild and domestic canids (definitive hosts: DHs), which in turn become infected by ingesting small rodents (intermediate hosts: IHs) carrying the larval stage (metacestode) of the pathogen [1]. *E. multilocularis* is the agent of a severe zoonosis, alveolar echinococcosis [2], which affects more than 18,000 new patients/year worldwide [3] and more than 150–200/year in the endemic area of central-eastern Europe.

In 2019, among the 751 human echinococcosis cases reported in the EU, 147 (26.5%) were attributable to *E. multilocularis* [4], with an incidence which has been increasing notably in recent decades [5]. In humans, who act as dead-end IHs, transmission is predominantly food-borne, and infection occurs when eggs (oncospheres) are ingested with water, wild berries and mushrooms or garden vegetables contaminated with DH faeces [6]. Oncospheres hatch in the gut, penetrate the intestinal wall and by the lympho-hematogenous route reach the liver, where they develop into metacestodes. These multiply asexually, infiltrate the liver and can spread to other organs through a metastasis-like process [7]. Alveolar echinococcosis develops very slowly in humans, taking from a few months [5] to 15 years (https://www.who.int/news-room/fact-sheets/detail/echinococcosis) to become clinically evident, although immune-suppressed patients show faster proliferation with earlier detection [8]. If untreated, prognosis is poor, and even the treatment itself is burdensome, as it includes a combination of surgery and long-term anti-parasitic therapy [9].

In the EU, the DH is predominantly represented by the red fox (*Vulpes vulpes*). From the historically endemic areas of Switzerland, southeast France and southern Germany, *E. multilocularis* has been expanding north, to northern France and Scandinavia, as well as east to the Balkans [4; 10-12]. Moreover, in the last two decades this cestode has been found sporadically in southern France [4; 13], as well as northern Italy in both the eastern [14] and western Alps [15], which represents at the moment the southernmost border of the parasite’s distribution in the EU. Although high spatial heterogeneity has been noted, in the endemic areas of central and northern Europe the host-parasite-environment pattern seems well established and is expected to be more predictable, whereas the same is not typical of the edges of the distribution [16]. In these areas, long-term surveillance is crucial to assess possible trends in prevalence and spread, as well as the exposure risk for humans. To this aim, sensitivity of the available diagnostic tests is of paramount importance, since worm burden levels are low and the presence of *E. multilocularis* infections in both DHs and IHs may be very hard to detect.

In the Province of Bolzano (Alto Adige, northeastern Alps, Italy), *E. multilocularis* in red foxes was first detected in early 2000 at a prevalence of about 13%, estimated using a nested PCR on DNA extracted from fox faeces [14]. In the following years, in order to increase the surveillance area and sample, another test (Coproscopy+Multiplex PCR [CMPCR] on parasite’s eggs) was used to analyse 2872 faecal samples across northeastern Italy. While optimizing the cost/benefit ratio [17], this test was known to have a good specificity (93.4%) but a low sensitivity (54.8%), relying on a preliminary screening by flotation of cestodes’ eggs, so that their burden could drop under the detection threshold [18]. CMPCR was used to confirm the persistence of the Alto Adige focus from 2012 to 2018, with an increasing trend in prevalence in later years, although never higher than 5% annually [17]. Considering the persistence and severity of this zoonosis, the current work aimed to enhance the sensitivity of *E. multilocularis* surveillance in red foxes, re-assess its prevalence in Alto Adige, and provide guidelines for a more effective surveillance strategy at the southern edge of its distribution.

## Methods

In 2019 and 2020, 235 red foxes that were legally culled or found as carcasses were collected across Alto Adige by Provincial wildlife technicians and transported to the Bolzano Laboratory of Istituto Zooprofilattico Sperimentale delle Venezie (IZSVe). To achieve an even sampling regime across the territory, one to four animals were collected from each hunting area (Fig 1). At necropsy, a faecal sample was taken from the rectum by a sterile glove. Moreover, when it was intact and not damaged (e.g. by the gun shot), the small intestine was tied off at both ends, removed and stored. Both faecal samples and small intestines were frozen at −80°C for at least 72h to inactivate *Echinococcus* eggs.Each faecal sample was then divided into two equal parts and tested for the presence of *E. multilocularis* by two methods:

**Fig 1.**
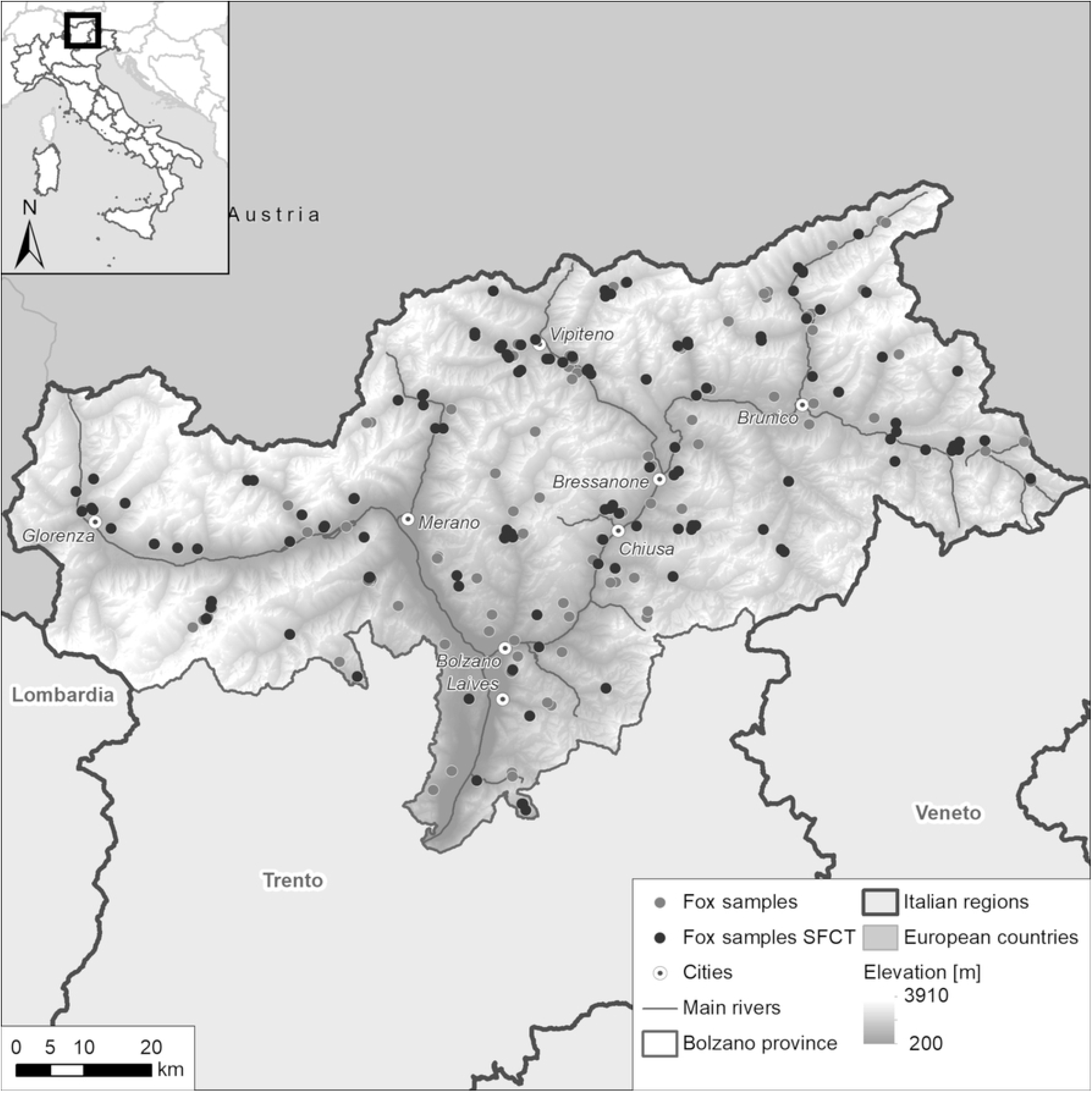
Distribution of individual fox faecal samples from Alto Adige in 2019-2020 (n=235) and subset examined for adult worms by SFCT (n=123)

1. CMPCR, following Citterio et al. 2021 [17] (at the Parasitology laboratory of IZSVe): briefly, 2 g of faecal matter was examined for *Taeniidae* eggs by flotation, filtration and sieving. Multiplex PCR amplification and Sanger sequencing were then performed on harvested eggs.
2. qPCR following Knapp et al. 2016 [19], with minor modifications (at the Animal Genetics laboratories and Sequencing Platform of the Fondazione E. Mach): in brief, whole DNA was extracted from 50 mg of faecal sample as described in Santa et al. 2019 [20], followed by qPCR amplification (Viia 7 Real-time PCR System, ThermoFisher Scientific, Waltham, MA) targeting the mitochondrial DNA marker rrnL, using 10 pmol of species-specific primers for *E. multilocularis* (202 bp) and 0.2 pmol of hydrolysis probe. The qPCR-positive samples were amplified with the same primer pair and sequenced with the dideoxy chain-termination protocol on an ABI PRISM 3730xl Genetic Analyzer (Applied Biosystems) using the BigDye Terminator cycle sequencing kit (Perkin Elmer, Applied Biosystems Division, Foster City, CA, USA). Sequences were aligned with an *E. multilocularis* reference fragment (GenBank acc. n. AB018440) using BioEdit 7.0.9 [21].

In order to evaluate the performances of the two diagnostic tests, a subsample of fox intestines, was examined for adult worms by the Scraping, Filtration and Counting Technique (SFCT), considered as the gold standard for *E. multilocularis* detection, following Santa et al. 2018 [22]. We performed SFCT on the available small intestines of the foxes that had tested positive for one or both diagnostic tests on faeces (n= 23), plus on a manageable number of fecal-negative samples (n=100), for a total of 123. Briefly, the pylorus to caecum section of the small intestine was cut into 30 cm lengths, each of which was then opened longitudinally and rinsed with tap water to remove loose faecal content. The intestine was then scraped, the resulting wash water was poured through a series of sieves (1000 μm, 212 μm and 75 μm), and the respective filtrates were collected separately into three distinct beakers. The filtrates were analysed by stereomicroscope (10X-63X) in 5 cm gridded petri dishes, to count the adult helminths. Using the SFCT results, sensitivity (Se) and specificity (Sp) with 95% exact binomial confidence intervals were calculated for each test, and their agreement (Cohen’s Kappa) to the standard was provided. True prevalence (TP) was then estimated, using the adjustment to the Rogan-Gladen formula as proposed by Lang and Reiczigel for confidence intervals [23]. Finally, the sample size needed for the detection of *E. multilocularis* by each of the two tests was assessed at the estimated TP and according to varying expected prevalence using the modified binomial approximation analysis method implemented in Epitools (https://epitools.ausvet.com.au/freecalctwo). We set a target population size of 300 individuals on a local scale (hypothetical epidemiological unit), based on the results of unpublished data from wildlife management offices in different areas of the northeastern Italian Alps, providing estimated densities of 2.1, 3.4 and 7.5 foxes/km^2^.

## Results

The distribution of the 235 fox faecal samples and of the 123 fox small intestines is shown in in Fig 1. Of the 235 fox faecal samples, seven (2.9%) were CMPCR-positive and 34 (14.4%) were qPCR-positive.

Adult *E. multilocularis* were found by SFCT in 24/123 (19.5%) fox small intestines (Fig 2) while, for the same animals, five (4.1%) and 23 (18.7%) faecal samples resulted as CMPCR and qPCR positive, respectively. Positive outcomes for the three diagnostics are summarized in Table 1.

**Table 1.** Matrix of the results of the two faecal tests (CMPCR and qPCR) and the gold standard (SFCT) for the detection of E. multilocularis in 123 foxes.

| Test | Coproscopy + Multiplex PCR (CMPCR) | qPCR | Scraping, filtration counting technique (SFCT) |
| --- | --- | --- | --- |
| CMPCR | 5 (4.1%) |  |  |
| qPCR | 5 | 23(18.7%) |  |
| SFCT | 5 | 20 | 24 (19.5%) |

**Fig 2.**
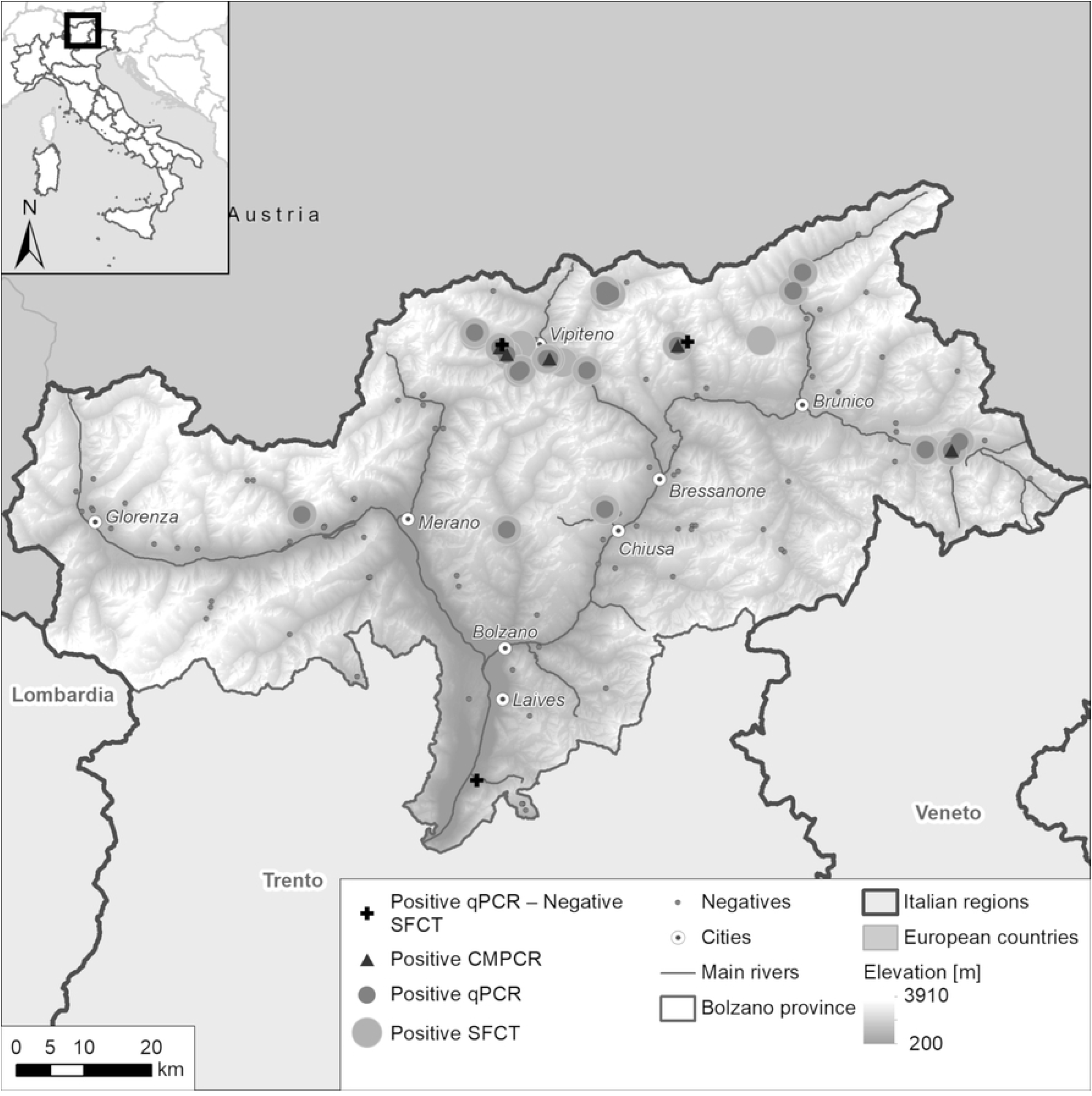
Results of the fox fecal samples collected in 2019-2020 (n=123) tested by Coproscopy+Multiplex PCR (CMPCR), qPCR and Scraping, Filtration and Counting Technique (SFCT-gold standard) for *E. multilocularis* in Alto Adige, Italy.

Compared to SFCT, CMPCR detected ‘true’ infections in 5/24 (4.1%) cases, resulting in very poor sensitivity (Se) (0.21; Table 2), and far lower than previously reported (about 0.55 [18]). Conversely, qPCR confirmed the SFCT results in 20/24 (86.9%) of the cases, translating into a much higher Se (0.83; Table 2). The qPCR test also identified three positive samples that were negative for SFCT, and thus considered as false positives.

**Table 2.**
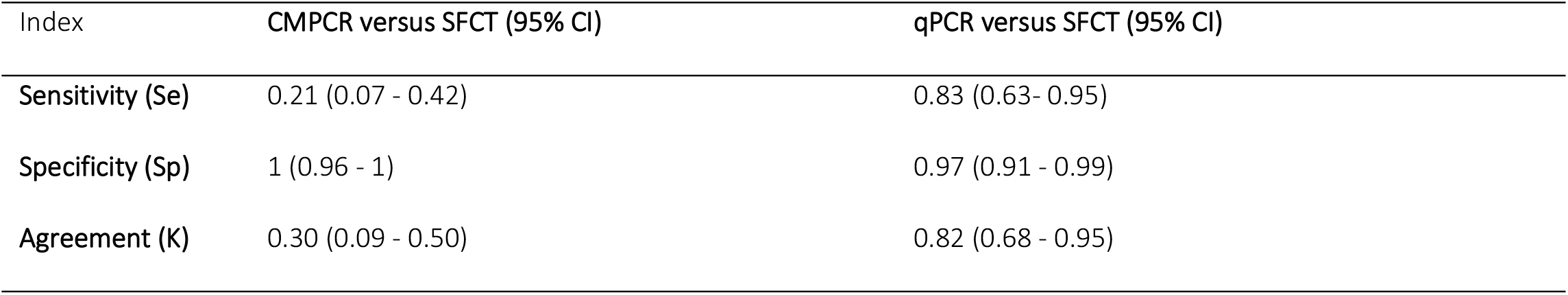
Sensitivity, specificity and agreement (K) for two diagnostic methods (Coproscopy+Multiplex PCR [CMPCR] and qPCR) for detecting E. multilocularis in fox fecal samples compared to the Scraping, Filtration and Counting Technique (SFCT - gold standard)

Based on the Se and Sp calculated from these results (Table 2), the TP estimated for the whole sample set (n=235) was 14.30 % (CI95%: 0-29.14) for CMPCR and 14.24% (CI95%: 7.04-22.64) for qPCR. The sample size required for the detection of *E. multilocularis* in a fox population of 300 individuals with various expected prevalences is shown in Fig 3. For example, if the expected prevalence is 15%, about 90 individuals (about one third of the population) should be tested when using CMPCR, whereas about 60 individuals (20%) would be enough to detect parasite presence by qPCR.

**Fig 3.**
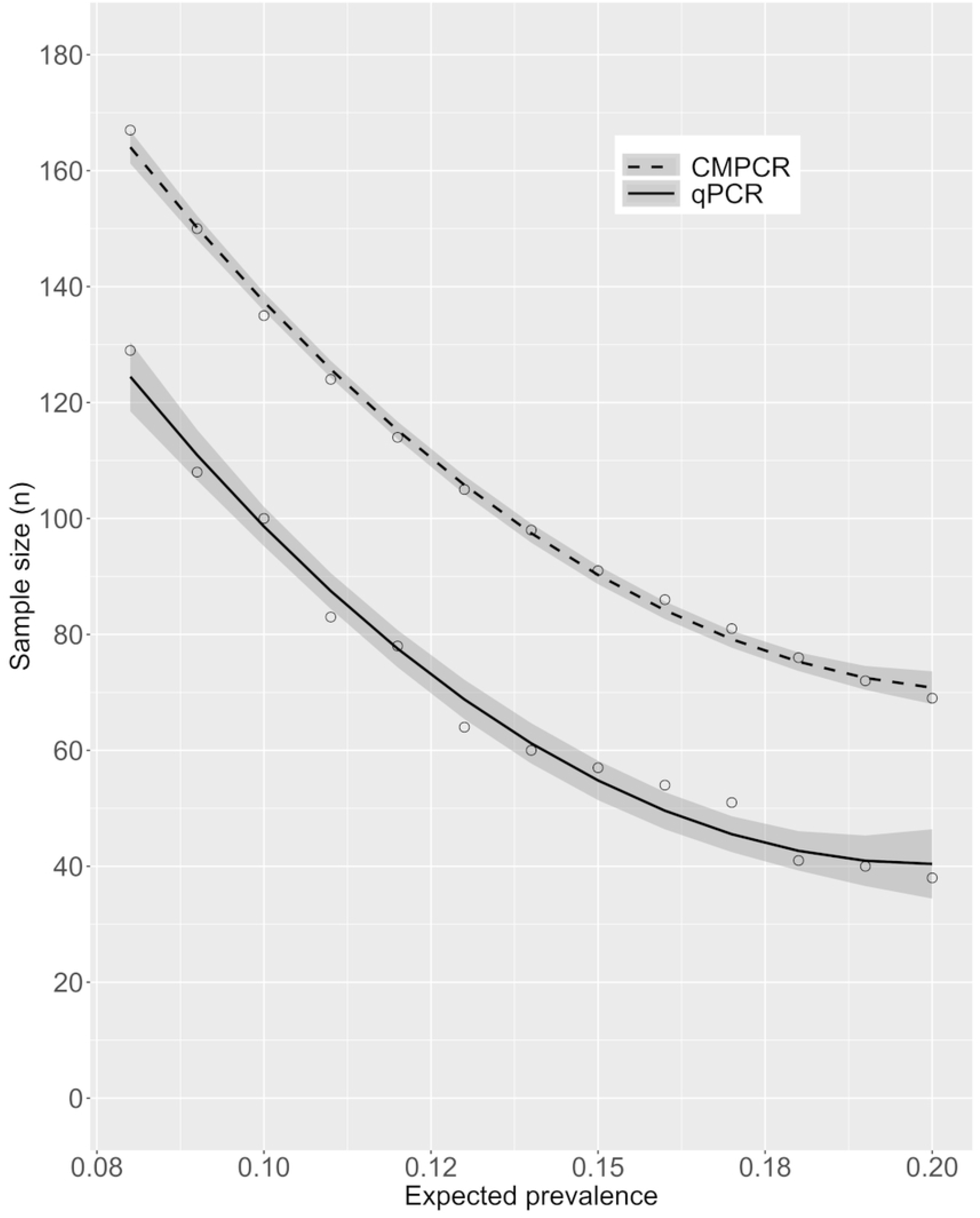
Sample size required to detect E. multilocularis at various expected prevalences (with 95% CI) in a population of 300 foxes, based on Sensitivity and Specificity of CMPCR and qPCR estimated here. Regression was obtained by interpolating 13 prevalence values.

## Discussion

Our results lead to the conclusion that the true prevalence of *E. multilocularis* in Alto Adige has been higher than previously reported in the last decade (<5%). According to the standards proposed by Casulli et al. (2015), this region should now be considered as highly endemic (prevalence >10%) [24], suggesting that surveillance should be intensified both locally and in the bordering territories.

The methods most widely used in the EU for detecting *E. multilocularis* in animal carcasses are modifications of techniques targeting adult parasites, such as the sedimentation and counting (SCT) and intestinal scraping (IST) [24]; the copro-antigen ELISA test [25]; or PCR [26] for species determination. However, methods aimed at detecting adult helminths have practical limitations, since they require the collection and necropsy of a consistent number of DHs throughout a study area, storage capacity of large volumes of organs temporarily at −80°C (for ensuring positive samples are not infective) and long-term at −20°C (for storage), as well as being time-consuming (at least 3 hours for each sample). Therefore, methods on faeces remain the best option for surveillance, given also that faecal samples can be collected in the field using a standardized protocol, and many samples can be screened simultaneously and rapidly. Surveillance coupling monitoring faeces of both fox carcasses and scats would better encompass the life cycle of both the parasite and its main definitive host throughout the year, and faecal prevalence could be used as a proxy for human risk of exposure. In addition, non-invasive collection and genotyping of scats would allow repeated sampling on the same fox social groups, tracking the seasonal pattern of infective eggs shedding at an individual level. Antigenic and molecular tests are rapid and useful in regions characterized by a high parasitic presence [27], but copro-antigen ELISA has the disadvantage of a low sensitivity, especially in the case of low parasitic burden [24]: molecular methods remain therefore the best option. Based on our results, tests directly applied on faeces, as qPCR, are recommended: in fact, besides reducing the number of samples needed to detect the infection (Fig 3) and consequently, the sampling effort and the cost of analyses, qPCR would also be the best option for detecting parasite DNA from scats collected in the field, since these generally result in poor quality DNA compared to fresh fecal samples taken during necropsies [28]. The higher sensitivity of qPCR is offset by an apparent decrease in specificity, suggested by the three positive samples not identified by SFCT (see also [29]). However, since SFCT is designed to detect adult worms, it may be that the extra positive samples detected by qPCR were indications of late infections, when traces of parasite DNA may still be present in the intestine but adult helminths are no longer macroscopically detectable or, on the contrary, of early infections, when immature parasites (scolices) are present but are very difficult to identify microscopically [14].

At the edges of *E. multilocularis* distribution, an efficient strategy for surveillance would be to progressively include areas bordering the foci, in which the parasite could have gone undetected, or in which *E. multilocularis* had been found occasionally, but no further confirmed, using qPCR as the main diagnostic tool. In such a framework, we suggest that surveillance in foxes should be performed on a small scale (e.g. considering an area similar to an Alpine valley as the basic epidemiological unit) rather than a larger one (e.g. using the NUTS at level 1 or 2, as recommended for *E. granulosus* by Tamarozzi et al. 2020 [30]). This would optimize efforts by increasing the chances of detecting *E. multilocularis* in areas previously considered free from the parasite. Moreover, such a detailed surveillance would be more informative of the actual risk of exposure for humans.

Finally, it is worth noting that a high endemicity of *E. multilocularis*, as that found in Alto Adige, should be a mandate for increased surveillance of possible human cases, as well as infections in domestic dogs. As dog infection could definitely increase the risk to humans, we suggest that this surveillance also use the most sensitive molecular tools available.

## Authors’ contributions

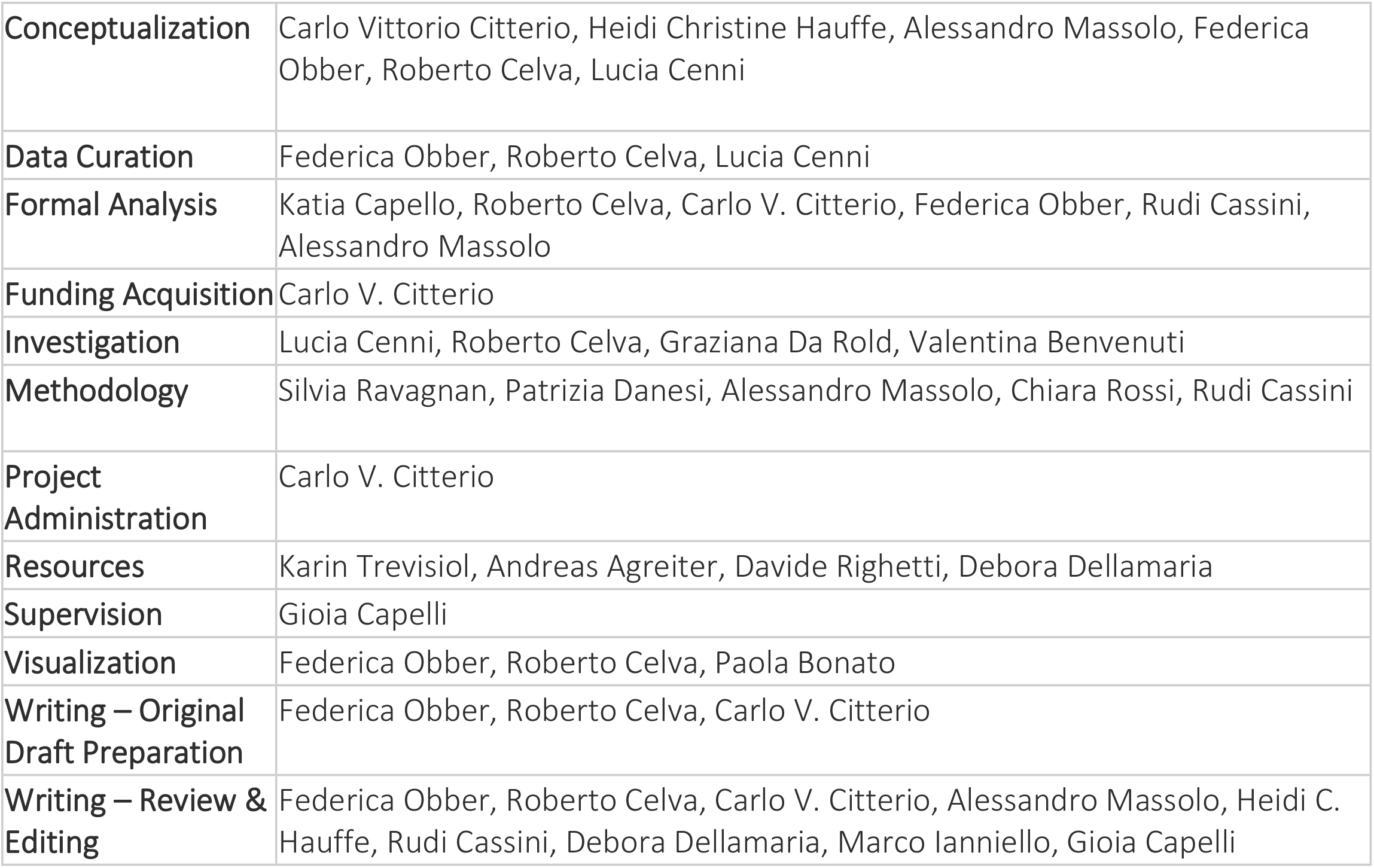

